# Integration of metabolic flux with hepatic glucagon signaling and gene expression profiles in the conscious dog

**DOI:** 10.1101/2023.09.28.559999

**Authors:** Katie C. Coate, Christopher J. Ramnanan, Marta Smith, Jason J. Winnick, Guillaume Kraft, Jose Irimia-Dominguez, Ben Farmer, E. Patrick Donahue, Peter J. Roach, Alan D. Cherrington, Dale S. Edgerton

**Author notes:** KCC and CJR are co-first authors. Deceased (Peter J. Roach). Dale Edgerton is the corresponding author.

## Abstract

Glucagon rapidly and profoundly simulates hepatic glucose production (HGP), but for reasons which are unclear, this effect normally wanes after a few hours, despite sustained plasma glucagon levels. This study characterized the time course and relevance (to metabolic flux) of glucagon-mediated molecular events in the livers of conscious dogs. Glucagon was either infused into the hepato-portal vein at a 6-fold basal rate in the presence of somatostatin and basal insulin, or it was maintained at a basal level in control studies. In one control group glucose remained at basal while in the other glucose was infused to match the hyperglycemia that occurred in the hyperglucagonemic group. Elevated glucagon caused a rapid (30 min) but only partially sustained increase in hepatic cAMP over 4h, a continued elevation in G6P, and activation and deactivation of glycogen phosphorylase and synthase activities, respectively. Net hepatic glycogenolysis and HGP increased rapidly, peaking at 30 min, then returned to baseline over the next three hours (although glucagon’s stimulatory effect on HGP was sustained relative to the hyperglycemic control group). Hepatic gluconeogenic flux did not increase due to lack of glucagon effect on substrate supply to the liver. Global gene expression profiling highlighted glucagon-regulated activation of genes involved in cellular respiration, metabolic processes, and signaling, and downregulation of genes involved in extracellular matrix assembly and development.

## Introduction

Stimulation of hepatic glucose production (HGP) by glucagon is critical to the regulation of glucose homeostasis during fasting (62), exercise (63), trauma and infection (53), and hypoglycemia (8, 47, 50). The physiologic effects of acute hyperglucagonemia in healthy animals and humans have been well-characterized. Glucagon stimulates glycogenolysis and causes a rapid and profound increase in HGP; an effect that characteristically wanes with time. Glucagon also plays a fundamental role in the pathogenesis of diabetes; increasing fasting glucose production and impairing hepatic glucose uptake (13, 46, 60, 61).

Early studies, primarily utilizing isolated liver slices and cell preparations, characterized glucagon’s effects on cAMP and enzymes in the glycogenolytic, gluconeogenic, and glycolytic pathways. More recent rodent studies have expanded our understanding of the molecular effects of hepatic glucagon signaling, describing the regulation of genes and proteins involved in glucose, lipid, and protein metabolism. Given the difficulty in carrying out trans-hepatic metabolic flux studies in the rodent, however, it has remained unclear how glucagon mediated effects on cellular signaling and hepatic gene expression correlate temporally with changes in metabolic flux. Moreover, the mechanisms underlying the waning of glucagon’s stimulatory effect on HGP are unclear.

Given these uncertainties, and the fact that glucagon is a potential target for the treatment of diabetes (4, 15, 46), examination of glucagon’s molecular regulation of metabolic flux in a large animal model is needed. Our primary aim was to characterize the time course of relevant glucagon-mediated cellular events during acute physiologic hyperglucagonemia and to correlate those changes with hepatic transcriptomic signatures and alterations in metabolic flux in the conscious dog.

## Methods and Materials

### Animal care and surgical procedures

Dogs of either sex were housed and fed as described previously (43, 44) and studied after an 18h fast. The surgical facility met the standards published by the American Association for the Accreditation of Laboratory Animal Care, and the protocols were approved by the Vanderbilt University Medical Center Animal Care Committee. Two weeks before experimentation, all dogs underwent a laparotomy in order to implant sampling catheters into the femoral artery, the hepatic portal vein, and the hepatic vein, as well as portal vein infusion catheters in the splenic and jujenal veins, and ultrasonic flow probes (Transonic Systems, Ithaca, NY) around the hepatic artery and the portal vein, as described previously (43, 44). All dogs were healthy (leukocyte count<18000/mm^3^, hematocrit>35%, good appetite, and normal stools).

### Experimental Design

Each study consisted of an equilibration (–150 to –30 min), a basal (–30 to 0 min) and an experimental period (30 min or 4h). At –150 min, [3-^3^H]glucose (priming dose of 35 µCi followed by 0.35 µCi/min) was administered intravenously (IV) and somatostatin was infused (0.8 µg/kg/min; Bachem, Torrance, CA) to inhibit the endocrine pancreas. Glucagon (0.57 ng/kg/min; Lilly, Indianapolis, IN) and insulin (as required to maintain euglycemia; Lilly) were infused intraportally at basal rates. The basal insulin infusion rate established in the equilibration period was maintained during the experimental period in each animal. Experiments were terminated at 4h in the basal glucagon/basal glucose control group (CTR) and at either 30 min or 4h in the basal glucagon/hyperglycemic group (GLU) and hyperglucagonemic/hyperglycemic (GGN+GLU) groups. In the CTR and GLU groups, basal intraportal glucagon infusion was maintained during the experimental period, whereas in the GGN+GLU group the rate was increased 6-fold (to 3.42 ng/kg/min). In the GLU group, intravenous glucose was infused to match the arterial plasma glucose level observed in the GGN+GLU group. Since CTR animals remained at a basal steady state, the 4h liver biopsy was assumed to provide reasonable control data for both the 30 min and 4h time points. Immediately after obtaining the final blood sample, each animal was anaesthetized with pentobarbital and a laparotomy was performed. The hormone and glucose infusions were continued while liver sections were rapidly freeze-clamped in situ and subsequently stored at –70°C, as described previously.

### Metabolite analysis

Hematocrit levels, glucose, insulin, and NEFA levels in plasma, and alanine, lactate, and glycerol concentrations in blood were determined using standard procedures as previously described (43). Plasma glucagon (GL-32K, Millipore-Sigma, Burlington, MA) was measured by radioimmunoassay. The Vanderbilt University Medical Center Analytical Services Core previously found that 54 ± 3% of what was measured by the GL-32K assay under baseline fasting conditions was nonspecific cross-reacting material rather than glucagon. In their analyses, glucagon was measured in plasma collected from five dogs before (37 ± 3 pg/ml; n=75 samples) and during somatostatin infusion (20 ± 1 pg/ml; i.e. the non-specific contribution; n=75). Complete suppression of endogenous glucagon secretion by somatostatin was verified by a total loss of the portal vein to arterial glucagon gradient (which was 11 ± 2 pg/ml before somatostatin infusion compared to 0 ± 1 pg/ml during it). Therefore, true glucagon levels were estimated by subtracting half of each dog’s basal glucagon level (i.e., the nonspecific component of the measurement) from measured values.

### Liver tissue analyses

Representative pieces of the 3 largest liver lobes were combined for analysis, SDS-PAGE and Western blotting was performed using standard methods (43). Fructose-2,6-bisphosphate (F_2,6_P_2_) concentration, glucose-6-phosphate (G6P) concentration, liver glycogen levels, and enzyme activities of glycogen synthase (GS) and glycogen phosphorylase (GP) were assessed using established methods (23, 36, 55, 59). cAMP was measured using the DetectX Cyclic AMP Chemiluminescent Direct Immunoassay kit (Arbor Assays, Ann Arbor, MI).

### Bulk RNA library preparation and sequencing

Bulk RNA sequencing (RNA-seq) was performed as described previously (22). Briefly, nucleic acid extraction and purification was performed on 9 canine liver biopsies collected under a steady-state conditions at the end of the 4h experimental period (n=3 each from CTR, GLU, and GGN+GLU groups). RNA concentration and integrity (RIN>7 for all samples) was assessed via Qubit fluorometric quantitation and tapestation bioanalyzer (Agilent), respectively. Libraries were prepared using the Takara SMART-Seq v4 Ultra low input RNA kit and analyzed on a NovaSeq platform (Illumina) using paired-end (PE100/150) reads with 40M total reads per sample. STAR aligner (v2.7.3a) was used for read alignment. Raw read counts were assessed using HTSeq (v0.11.2) and normalized using DESeq2. Aligned reads were used for estimating expression of the genes using cufflinks (v2.2.1). Differentially expressed genes (DEGs) were determined using DESeq2 (R Bioconductor package, version 3.11) on the basis of fold change (cutoff ≥ ±1.0) and adjusted p-value (<0.1) as determined by the Benjamini-Hochberg false discovery rate method (5). To delineate glucagon-regulated pathways and biological processes, significantly enriched groups of coordinately regulated genes were identified by Gene Set Enrichment Analysis (GSEA) using Hallmark genesets (30) and gene ontology (GO) terms from the Molecular Signatures Database (56).

### Calculations

Net hepatic glucose balance was assessed using the arteriovenous difference technique as described previously (43, 44). Gluconeogenic flux was determined by taking the sum of net hepatic uptake rates of gluconeogenic precursors and dividing by two to account for three carbon precursor incorporation into one six-carbon glucose molecule. Glycolytic flux was estimated by taking the sum of net hepatic output rates (when present) of the substrates noted above (in glucose equivalents) and hepatic glucose oxidation (assumed to be 0.2±0.1 mg/kg/min as previously validated) (38, 51). Glucose turnover, used to determine glucose production, was determined using the GLUTRAN circulatory model of Mari et al (35).

### Statistical analysis

Statistical comparisons were carried out using one-way ANOVA (Prism10) or two-way repeated measures ANOVA (group x time) (SigmaStat) with appropriate post hoc tests when significant F ratios were obtained. Significance was determined as P<0.05.

## Results

### Hormone levels

Insulin was maintained at a basal level in all groups (Fig. 1A, B). Arterial and hepatic portal vein glucagon levels were clamped at basal values in the CTR and GLU groups or increased to approximately 100 and 250 pg/ml, respectively, in the GGN+GLU group during the experimental period (Fig. 1C, D). There was about a 6-fold rise in glucagon in both vessels, in keeping with the portal vein glucagon infusion rate.

**Fig. 1.**
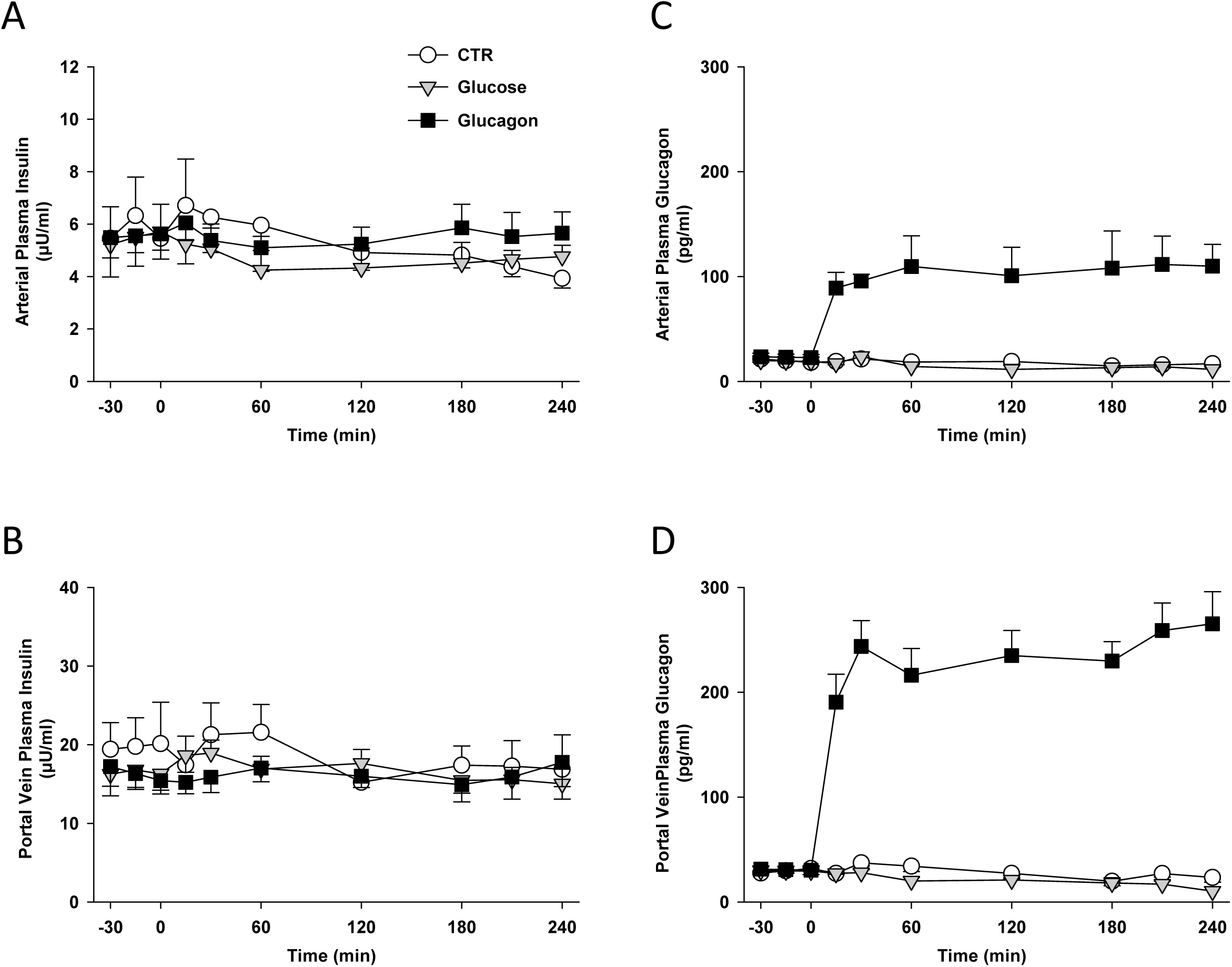
Hormone levels. (A) Arterial plasma insulin, (B) hepatic portal vein plasma insulin, (C) arterial plasma glucagon, and (D) hepatic portal vein plasma glucagon concentrations in 18h fasted conscious dogs during the basal (–30 to 0 min) and experimental (0-240 min) periods. Data are means ± SEM, in the CTR (n=6 from –30 to 240 min), GLU (n=8 from –30 to 30 min; n=4 from 30 to 240 min), and GGN+GLU (n=8 from –30 to 30 min; n=4 from 30 to 240 min) groups. Arterial and portal vein plasma glucagon levels were greater (*P*<0.05) in the GGN+GLU group compared to the GLU group between 15-240 min.

### Glucose, glycogenolytic, and gluconeogenic flux rates

Euglycemia was maintained in the CTR group, whereas hyperglycemia rapidly developed due to the effects of glucagon in the GGN+GLU group (Fig. 2A). This glycemic rise was matched in the GLU group using IV glucose infusion (Fig. 2B). During the basal period, the rates of net hepatic glucose balance (NHGB; Fig. 2C) and hepatic glucose production (Fig. 2D) were similar between groups. Hyperglucagonemia markedly stimulated both NHGB (5-fold) and HGP (4-fold) within 15 min (Fig. 2C, D), after which glucose production gradually waned and returned to baseline by the fourth hour of the clamp. When glucagon-stimulated hyperglycemia was matched by glucose infusion in the GLU group, there was a rapid decline in NHGB that resulted in a switch to slight net hepatic glucose uptake (∼1 mg/kg/min) by the end of the experiment (Fig. 3C). Consistent with this observation, hyperglycemia almost completely suppressed HGP (Fig. 2D). Therefore, although glucagon’s effect at 4h was reduced by 70% from its peak, it continued to effectively stimulate glucose production relative to that in the GLU group.

**Fig. 2.**
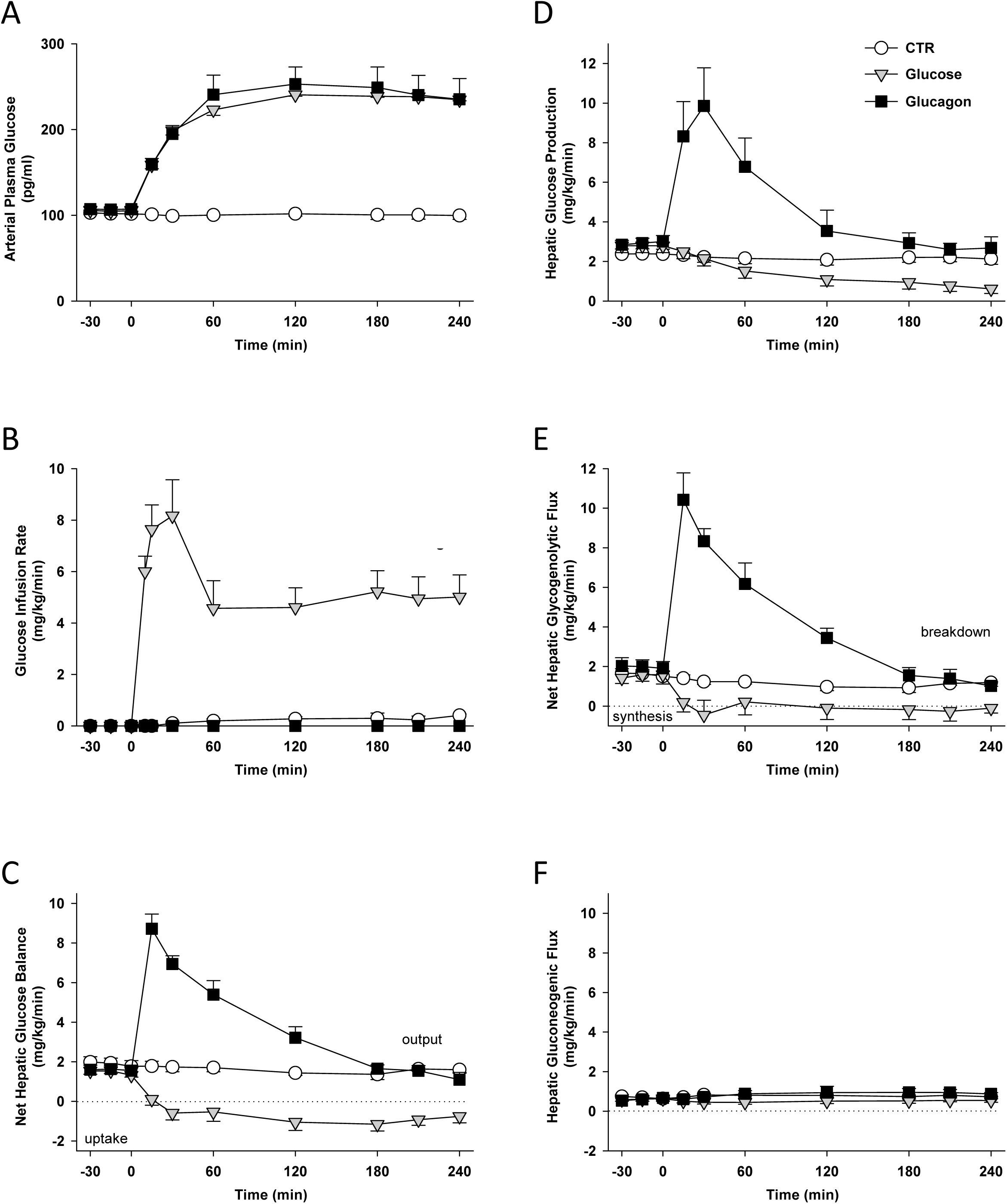
Glucose metabolism. (A) Arterial plasma glucose levels, (B) glucose infusion rates, (C) net hepatic glucose balance, (D) glucose production, (E) net hepatic glycogenolytic flux, and (F) hepatic gluconeogenic flux in 18h fasted conscious dogs during the basal (–30 to 0 min) and experimental (0-240 min) periods. Data are means ± SEM, in the CTR (n=6 from –30 to 240 min), GLU (n=8 from –30 to 30 min; n=4 from 30 to 240 min), and GGN+GLU (n=8 from –30 to 30 min; n=4 from 30 to 240 min) groups. Arterial plasma glucose levels differed (*P* <0.05) in the GGN+GLU and GLU groups compared to the CTR group between 15-240 min. The glucose infusion rate differed (*P* <0.05) in the GLU group compared to the other groups between 15-240 min. Net hepatic glucose balance, glucose production, and net hepatic glycogenolysis were greater (*P* <0.05) in the GGN+GLU group compared to the GLU group between 15-240 min.

**Fig. 3.**
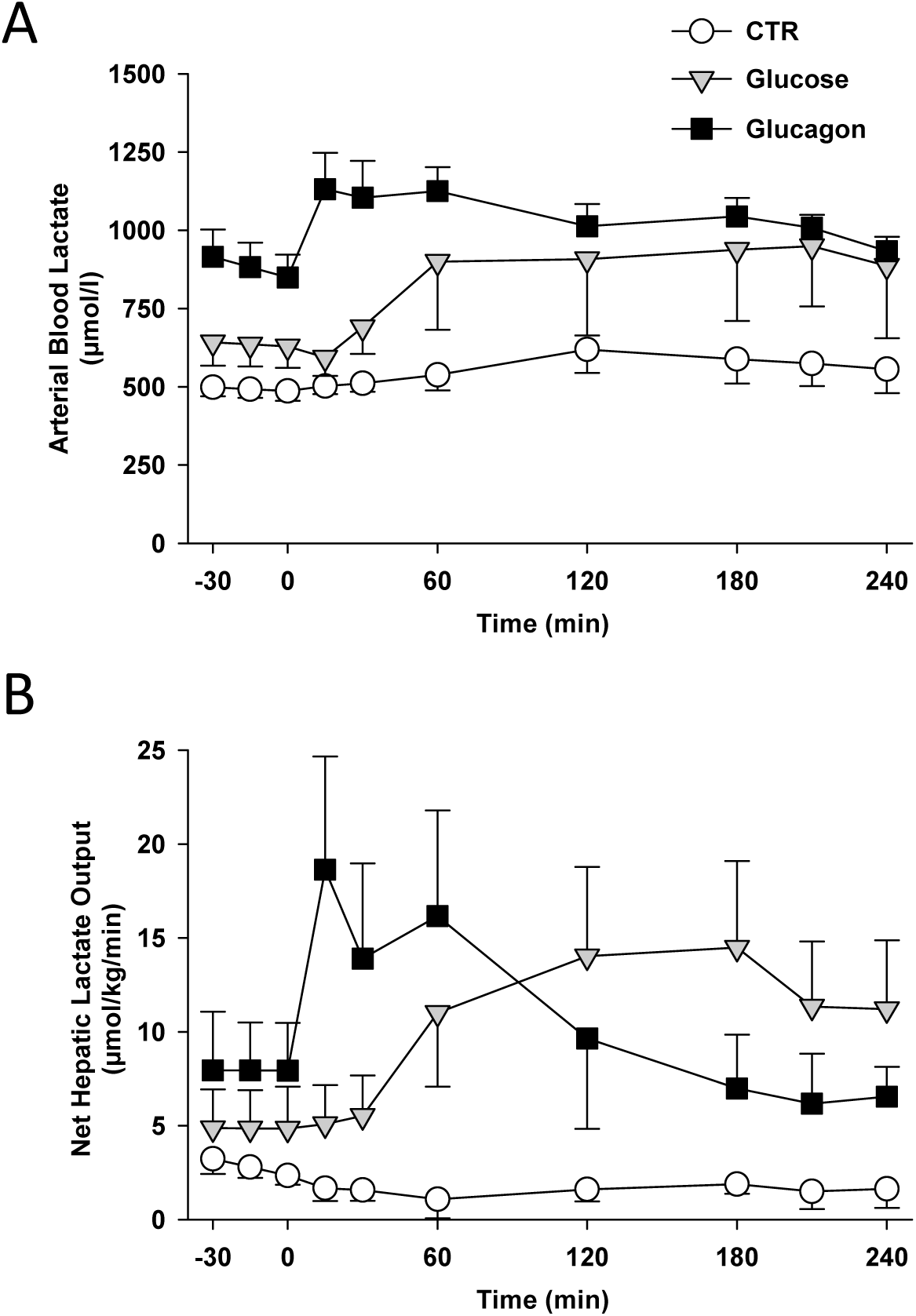
Lactate metabolism. (A) Arterial blood lactate and (B) net hepatic lactate production in 18h fasted conscious dogs during the basal (–30 to 0 min) and experimental (0-240 min) periods. Data are means ± SEM, in the CTR (n=6 from –30 to 240 min), GLU (n=8 from –30 to 30 min; n=4 from 30 to 240 min), and GGN+GLU (n=8 from –30 to 30 min; n=4 from 30 to 240 min) groups. Net hepatic lactate output was less (*P* <0.05) in the GGN+GLU group compared to the GLU group between 180-240 min, and it was greater in the GLU group than the CTR group throughout the study.

The rapid effects of glucagon (stimulatory) and glucose (suppressive) on liver glucose flux were both attributable to changes in net hepatic glycogenolytic flux (Fig. 2E). By contrast, gluconeogenic flux was not significantly altered by either treatment (Fig. 2F). The fact that HGP was reduced by hyperglycemia in the GLU group whereas hepatic gluconeogenic flux (to G6P) was not indicates that the gluconeogenic carbon was being used to form liver glycogen rather than plasma glucose. This is in line with the switch towards net hepatic glycogen synthesis seen in the GLU group (Fig. 2E).

### Lactate, fat, and amino acid metabolism

Arterial blood lactate levels were not altered over time in the CTR group, but they tended to increase in the GLU and GGN+GLU groups (Fig. 3A). Net hepatic lactate output, one indicator of hepatic glycolysis, remained at a basal rate throughout in CTR animals but increased after a delay of 30 min in GLU and then remained elevated (Fig. 3B). Glucagon, on the other hand, caused a rapid (within 15 min) stimulation of net hepatic lactate output, followed by a steady decline such that by the end of the study lactate output was below baseline. Thus, glucagon caused an initial surge in glycolytic flux but after 1h, limited the stimulatory effect of hyperglycemia on this pathway. Arterial levels and net hepatic balances of NEFA and glycerol did not differ between groups, nor did they change over time, suggesting a lack of effect of glucagon or glucose on lipolysis or NEFA reesterification under these conditions (Fig. S1). In the CTR and GLU groups there were no significant changes in arterial blood alanine levels (a representative gluconeogenic amino acid), net hepatic uptake, or net hepatic fractional extraction (Fig. S2). On the other hand, hyperglucagonemia increased hepatic alanine fractional extraction and net hepatic alanine uptake which caused a decrease in blood alanine levels.

### Cellular effects of a physiologic increase in glucagon

*Insulin signaling*. In the presence of basal insulin, elevations in glucagon or glucose had little effect on Akt phosphorylation. Glucagon abrogated glucose stimulated increases in GSK-3β and FOXO1 phosphorylation by 4h (Fig. 4A, 4B, 4C). *AMPK signaling*. Neither hyperglycemia nor hyperglucagonemia had consistent effects on the phosphorylation of AMPK at Thr172 or Ser485, or on ACC phosphorylation (an index of active AMPK) (Fig. 4D, 4E, 4F).

**Fig. 4.**
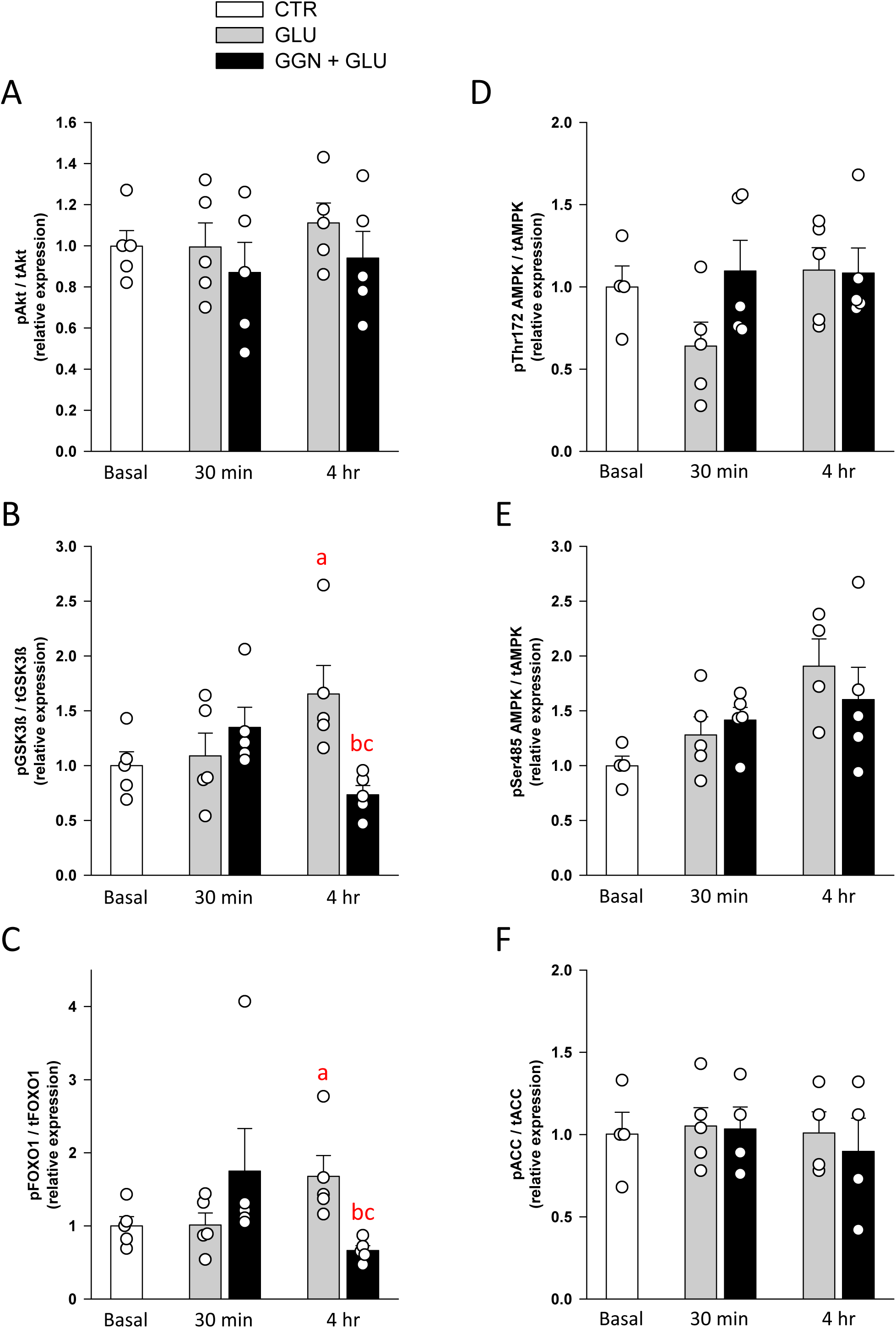
Phosphorylation state of (A-C) hepatic insulin signaling and (D-F) AMPK signaling proteins. Hepatic (A) Akt (p/t), (B) GSK3β (p/t), (C) FOXO1 (p/t), (D) AMPK (Thr172 p/t), (E) AMPK (Ser485 p/t), and (F) ACC (p/t). See Supplemental Table 1 for representative Western blots. Data are means ± SEM in the Control (n=4-5), Glucose (n=4-5), and Glucagon (n=4-5) groups for each time point. ^a^*P* <0.05 GLU or GGN+GLU vs CTR, ^b^*P* <0.05 GLU vs GGN+GLU within the same time point (30 min or 4h), ^c^*P* <0.05 GLU vs GLU or GGN+GLU vs GGN+GLU between time points (30 min vs 4h).

*Regulators of glycogen metabolism*. While the hepatic cAMP level was not altered by hyperglycemia, it was markedly elevated in the GGN+GLU group at 30 min (7-fold) and remained elevated at 4h, albeit at a somewhat lower level (4-fold; Fig. 5A). cAMP ultimately determines the extent of phosphorylation of glycogen phosphorylase (GP) and glycogen synthase (GS), thus, as expected, GP activity was markedly elevated at both 30 min and 4h in the GGN+GLU group (Fig. 5B). Although GP activity waned modestly over time (∼25% lower at 4h than 30 min), this also tended to occur in the GLU group (Fig. 5B), suggesting that the small decrease in activity may have been glucose-induced and that glucagon’s effect on phosphorylase activity was for the most part sustained. GS activity was not altered by hyperglycemia but decreased progressively in response to hyperglucagonemia (Fig. 5C). The ratio of GP to GS activities (an index of the net change in glycogen enzyme activity) was 12±2 in the CTR group. In the GLU group this ratio was 13±5 at 30 min and 8±1 at 4h. The ratios were 40±2 and 52±9, respectively, in the GGN+GLU group. Thus, in response to glucagon, the activity of the glycogen metabolizing enzymes shifted towards breakdown, and there is no evidence that a change in the phosphorylation of these enzymes explains why glucose production waned over time. It should be noted, however, that liver GS is hyperphosphorylated, and a small decrease in measured (*in vitro*) hepatic GS activity would not necessarily have a major effect on actual GS flux (the assay measures phosphorylation-dependent rather than allosterically regulated GS activity).

**Fig. 5.**
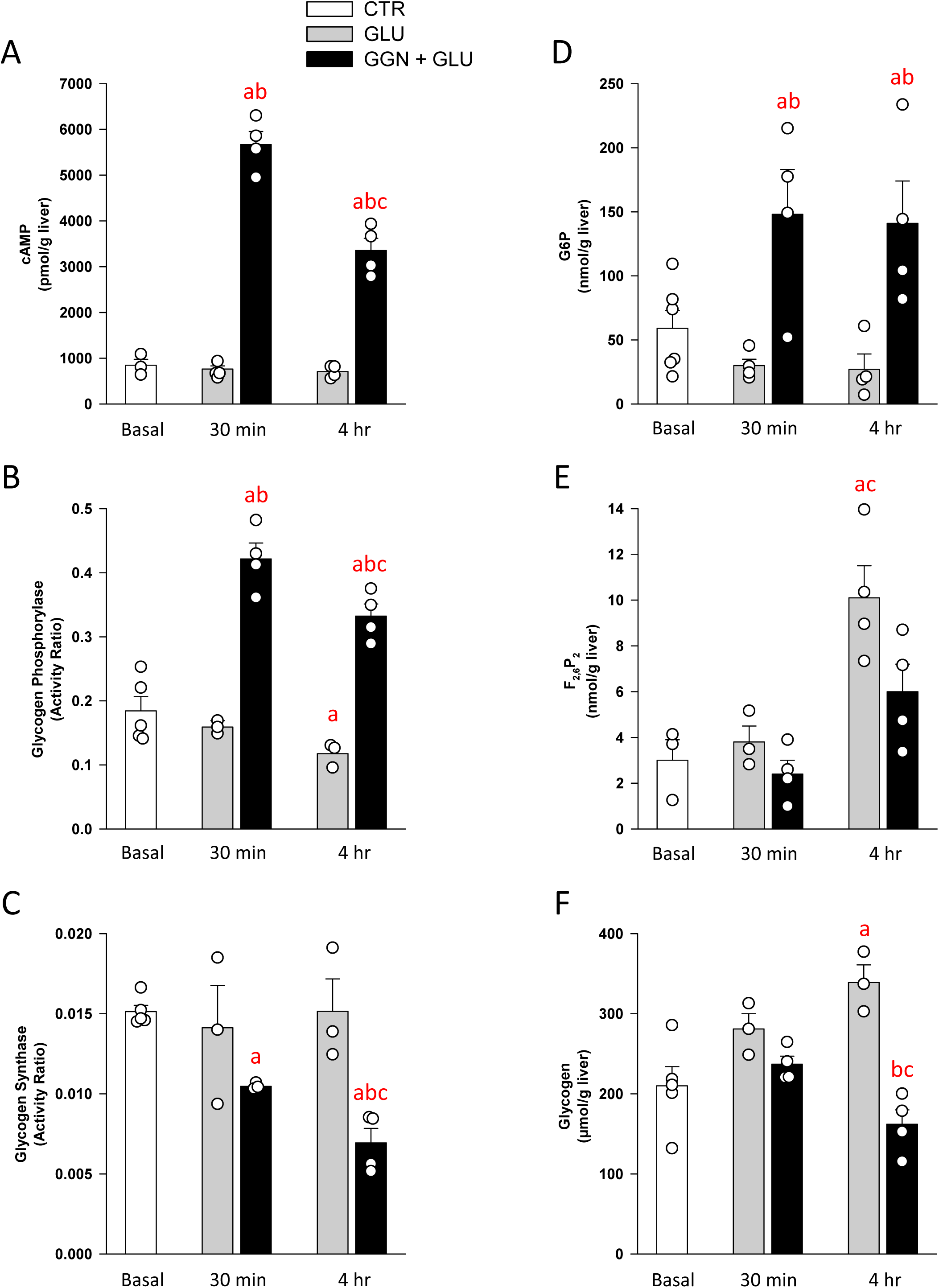
Hepatic cAMP and regulators of hepatic glycogen metabolism and glycolysis. Hepatic (A) cAMP, (B) glycogen phosphorylase activity ratio, (C) glycogen synthase activity ratio, (D) G6P, (E) F_2,6_P_2_, and (F) glycogen, Data are means ± SEM in the CTR (n=3-4), GLU (n=3-4), and GGN+GLU (n=4) groups for each time point. ^a^*P* <0.05 GLU or GGN+GLU vs CTR, ^b^*P* <0.05 GLU vs GGN+GLU within the same time point (30 min or 4h), ^c^*P* <0.05 GLU vs GLU or GGN+GLU vs GGN+GLU between time points (30 min vs 4h).

*Hepatic G6P, F_2,6_P_2_*, and *glycogen.* While hyperglycemia tended to reduce hepatic G6P content, hyperglucagonemia elicited a rapid (within 30 min), sustained (up to 4h), and marked increase in G6P (Fig. 5D). Hepatic GS is primarily regulated by the allosteric effects of G6P, thus it is likely that GS flux was enhanced in the GGN+GLU group (despite decreased *in vitro* GS activity; Fig 5C), at least in some hepatocytes (27), although this was clearly offset by a greater increase in GP flux during the early response to glucagon (Fig. 2E). Later there was a shift in net GS/GP balance which was less favorable to net degradation, likely due to prolonged and sustained elevations of G6P and glucose. Hepatic levels of F_2,6_P_2_, an allosteric activator of glycolysis and inhibitor of gluconeogenic flux, was markedly (3-fold) increased by hyperglycemia at 4h, whereas glucagon limited this glucose induced rise by 60% (Fig. 5E). Consistent with the cessation of net hepatic glycogen breakdown in the GLU group, glycogen levels increased progressively over time relative to the other groups (Fig. 5F). As expected given the high rate of net hepatic glycogenolysis in the GGN+GLU group, glucagon reduced the liver glycogen content (by more than 50% compared to the GLU group at 4h).

### Glucagon-regulated hepatic transcriptional signatures identified by bulk RNA-seq

To determine the hepatic transcriptional profiles uniquely associated with a physiologic increase in glucagon (GGN+GLU) compared with matched hyperglycemia (GLU) and baseline controls (CTR), we performed bulk RNA-seq of liver biopsies obtained from the GGN+GLU, GLU, and CTR groups at the end of the 4h experimental period. Comparative analysis of gene expression profiles in the GGN+GLU vs. GLU group revealed a total of 559 differentially expressed protein-coding genes (DEGs) of which 318 were up-regulated by hyperglucagonemia and 241 were downregulated (Fig. 6A). As anticipated, genes associated with gluconeogenesis (*G6PC* and *PPARGC1A*), amino acid transport and metabolism (*SLC1A2*, *SLC7A2*, *OAT* and *TAT*), and cAMP signaling (*ADCY1* and *PDE4B*) were among the most highly upregulated genes in the GGN+GLU group. Interestingly, the gene encoding follistatin (i.e., *FST*), a glucagon– and FOXO1-regulated hepatokine that promotes insulin resistance, white adipose tissue lipolysis, and hyperglycemia in rodents (57, 58), was likewise one of the most significantly induced genes by hyperglucagonemia. On the other hand, genes associated with cellular responses to carbohydrate stimuli (*PRKCE*, *KCNB1*, *EGR1*, and *FGF21*) as well as the matrisome (i.e., extracellular matrix [ECM] related, including *THBS1*, *MMP11*, *COL3A1*, *COL6A1*, and *INHBA*) were among the most significantly downregulated genes in the GGN+GLU group. Of note, the waning of net hepatic glycogenolytic flux in the GGN+GLU group correlated with marked downregulation of *MGAM2*, a gene that encodes maltase-glucoamylase 2, an enzyme predicted to be involved in glycogen breakdown through its alpha-1,4-glucosidase activity (Fig. 6A).

**Fig. 6.**
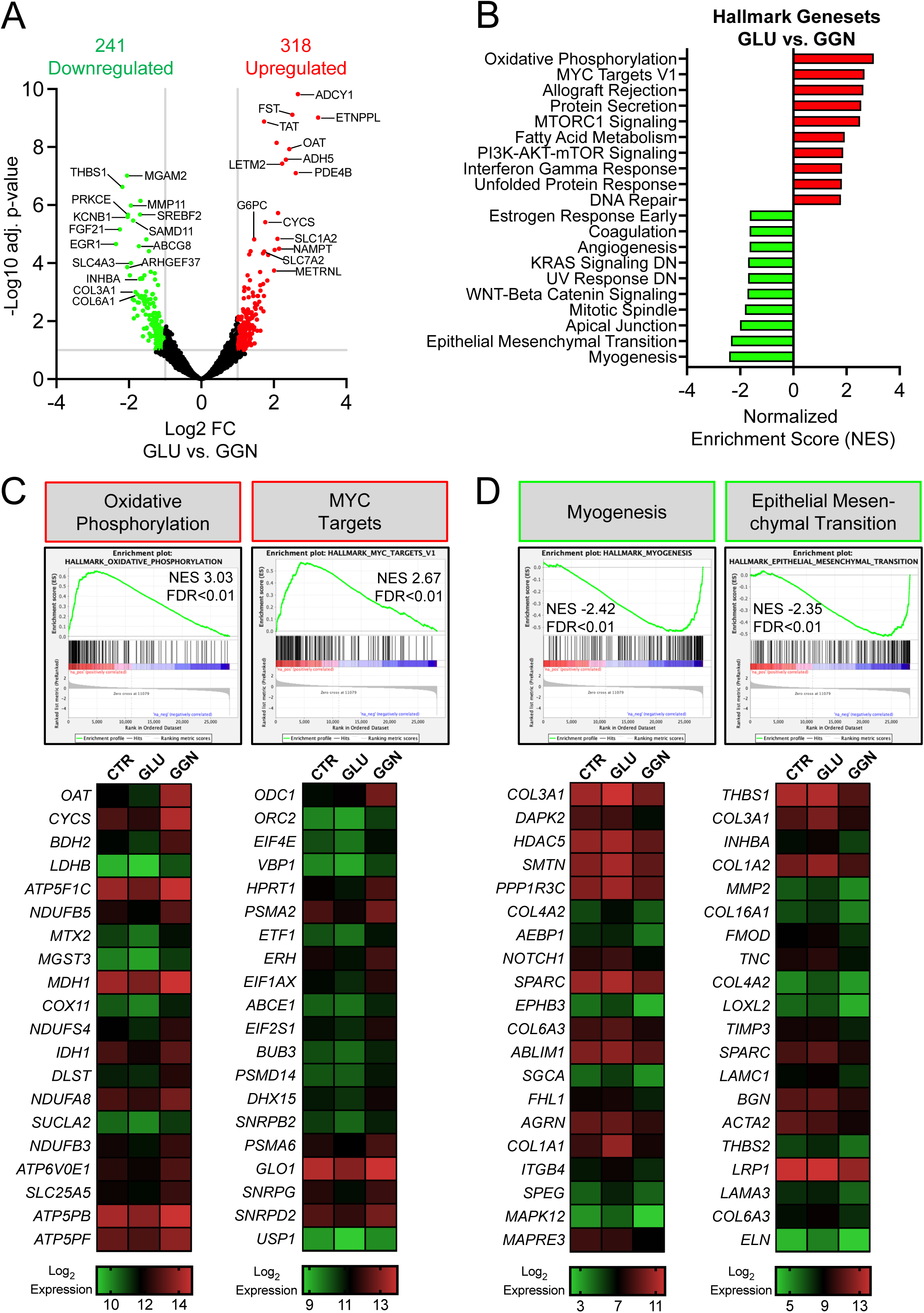
Glucagon-regulated hepatic transcriptional profiles identified by bulk RNA-seq. (A) Volcano plot illustrating the log_2_ fold change (FC; x-axis) and adjusted (adj.) p-values (y-axis) of all differentially expressed genes (DEGs) in GLU vs. GGN+GLU. A total of 559 protein-coding genes showed significant differential expression (i.e., –log10 adj. p-value≥1.0 and log_2_FC≥±1.0, as denoted by grey lines) in GLU vs. GGN+GLU: 318 up-regulated (red circles) and 241 down-regulated (green circles). **(B)** The top 10 significantly (FDR q-value < 0.1) enriched Hallmark genesets in GLU vs. GGN+GLU as determined by Gene Set Enrichment Analysis (GSEA), with the x-axis displaying normalized enrichment scores (NES) for positively (red bars; up-regulated genes in the GGN+GLU group are enriched in these genesets) and negatively (green bars; down-regulated genes in the GGN+GLU group are enriched in these genesets). **(C and D)** Heat map representation of the top 20 leading-edge transcripts underlying the **(C)** positive enrichment signals for oxidative phosphorylation (left panel) and MYC targets (right panel) or the **(D)** negative enrichment signals for myogenesis (left panel) and epithelial mesenchymal transition (right panel) as identified in panel B. Each row represents a different glucagon-regulated gene, whereas each column represents the mean log_2_ normalized expression value for these genes in the CTR, GLU, and GGN+GLU groups (n=3 each). Expression values are represented as colors and range from green (lowest expression) to red (highest expression).

### Gene set enrichment analysis (GSEA)

To identify pathways and biological processes significantly enriched in GGN+GLU vs. GLU groups, we performed GSEA using Hallmark genesets and GO terms from the MSigDB (30, 56). A total of 37 (out of 50) Hallmark genesets were significantly enriched (FDR<0.1) in the GGN+GLU group, with 24 positively and 13 negatively enriched (Fig. 6B). The most positively enriched glucagon-regulated pathway highlighted widespread induction of genes involved in hepatic mitochondrial oxidative phosphorylation (OXPHOS) and contained significant enrichment of related functional GO terms such as mitochondrial respiratory chain complex assembly, ATP synthesis coupled electron transport, NADH dehydrogenase complex assembly, mitochondrial electron transport chain NADH to ubiquinone, mitochondrial translation, and ribosome biogenesis. Leading-edge genes underlying the glucagon-regulated OXPHOS signal included those encoding mitochondrial respiratory chain complex proteins and ATP synthesis/transport proteins, such as *CYCS, COX11, NDUFB5, NDUFS4, NDUFA8, NDUFB3, ATP5F1C, ATP6V0E1, ATP5PB, ATP5PF,* and *SLC25A5* (Fig. 6C). Genes annotated to the MYC Targets geneset were also significantly enriched among the up-regulated genes with hyperglucagonemia, suggestive of heightened MYC transcription factor activation in the GGN+GLU vs. GLU group (Fig. 6B, 6C). Indeed, elevated cAMP-PKA signaling was shown to augment MYC protein expression *in vitro* (9), and MYC directly regulates genes that promote mitochondrial biogenesis and oxidative metabolism (39). In support of this, GO terms associated with leading edge genes underlying positive enrichment of the MYC Targets geneset included ribonucleoprotein complex biogenesis, translational initiation, peptide biosynthetic processes, spliceosomal complex assembly, and RNA binding. These findings suggest that MYC may be a molecular effector of glucagon-regulated mitochondrial OXPHOS gene expression in the liver. Protein secretion, mTORC1 signaling, and fatty acid metabolism were among the other positively enriched pathways in the GGN+GLU vs. GLU group (Fig. 6B and Fig. S3).

By contrast, genes annotated to pathways more broadly associated with regulation of ECM organization and interactions, including myogenesis, epithelial mesenchymal transition (EMT), and apical junction were significantly enriched among the downregulated genes with hyperglucagonemia (e.g., *COL3A1*, *COL4A2*, *COL6A3*, *COL1A2*, *THBS1*, *MMP2*, *TNC*, *FMOD*, *INHBA*) (Fig. 6B, 6D).

Likewise, GO terms related to collagen fibril organization, external encapsulating structure organization, ECM assembly, semaphoring plexin signaling pathway, and aorta development and morphogenesis were overrepresented among the downregulated genes with hyperglucagonemia. Pathological elevation in the expression of leading-edge genes underlying the signals in myogenesis and EMT, for example, have been shown to occur in response to liver injury and/or obesity and precede the development of hepatic fibrosis and insulin resistance (40, 64). Additionally, T2D in humans is associated with hepatic and extra-hepatic ECM remodeling (40, 64). Our data suggest that in healthy dogs, acute (i.e., 4h) hyperglucagonemia limits hepatic ECM proliferation and/or remodeling via transcriptional downregulation of genes involved in these processes. Whether these changes impact the acute stimulatory effect of glucagon on hepatic glucose production and/or amino acid metabolism remains to be determined.

## Discussion

A physiologic 6-fold rise in glucagon rapidly stimulated HGP and caused a rise in glycemia. Glucagon’s effect on the liver was transient, peaking within 15-30 min, with HGP returning to basal by 3h (although hyperglucagonemia continued to stimulate glucose production since matching hyperglycemia suppressed HGP below basal). The stimulation and waning of glucose production in response to glucagon was entirely the result of alterations in glycogen flux, with no detectable modification of hepatic gluconeogenic flux. The purpose of this study was to identify cellular and molecular correlates for the hepatic metabolic response to glucagon.

Since a selective rise in glucagon (insulin kept basal) causes hyperglycemia, and hyperglycemia has independent effects on liver glucose metabolism, the effects of hyperglycemia must be considered when evaluating glucagon action. While the ability of glucagon to stimulate HGP has been shown to be independent of the accompanying glycemic state (10, 12, 52), hyperglycemia has its own independent inhibitory effect on HGP. Thus, we included a matched hyperglycemic control group to isolate the unique metabolic and molecular effects of glucagon per se. Within 15 min hyperglycemia per se, in the presence of basal glucagon and insulin, completely inhibited net hepatic glucose balance without yet affecting HGP, indicating that hepatic glucose uptake rapidly increased while glucose cycling initially allowed hepatic glucose release to continue. At this time net hepatic glycogenolysis was suppressed to near zero, there was no change in glycolytic flux, and gluconeogenic flux persisted, indicating that gluconeogenesis was almost completely responsible for hepatic glucose release. By 30 min the liver had switched to low rates of net glucose uptake and net glycogen synthesis in the GLU group, HGP was reduced by more than 20%, and an increase (34%) in liver glycogen was detected. We did not detect any significant impact of hyperglycemia on markers of insulin signaling or regulators of glycogen metabolism or gluconeogenesis at this time. While changes in GS and GP activities in response to hyperglycemia were not evident, it should be noted that these assays reflect only the covalent state of the enzymes, and do not account for the impact of allosteric effectors on their overall activities (48). Therefore, an *in vivo* change in the activity of these enzymes may have gone undetected in our *in vitro* assay. At 30 min, net glycogen flux decreased to near zero, and G6P content was about 50% of basal. While a fall in G6P would be expected to shift flux towards net glycogen breakdown, hyperglycemia can cause the arrest of glycogenolysis and an increase in glycogen synthesis (3, 7). By 60 min, hyperglycemia had suppressed HGP by 50%, and glycolytic flux began to increase (as indicated by the increase in net hepatic lactate output). Indeed, F_2,6_P_2_ levels were elevated by the end of the study, which would be expected to promote glycolytic flux by increasing the activity of phosphofructokinase 1 (PFK-1). Whereas hyperglycemia did not eliminate hepatic gluconeogenic flux (conversion of non-carbohydrate precursors into G6P), gluconeogenesis per se (glucose derived from non-carbohydrate precursors that is released into the blood) was reduced, since HGP was nearly completely suppressed after 4 h. This suggests that gluconeogenic carbon was redirected from release as glucose to other fates. In support of this liver lactate output increased and glycogen levels were higher in the GLU group by the end of the study.

In contrast, glucagon stimulated a rapid (within 15 min) and profound increase in net hepatic glycogenolysis that was dominant over the suppressive effect of hyperglycemia. This glycogenolytic response was most likely caused by increased hepatic cAMP and its effects on GP and GS activities, which overrode any opposing allosteric effects of G6P or glucose on their activities. Glycogen breakdown provided substrate for increased glucose production and glycolytic flux, which were both tightly correlated with net hepatic glycogenolysis throughout the study. F_2,6_P_2_ is known to be increased by glucose and decreased by glucagon and to allosterically inhibit gluconeogenic flux while promoting glycolytic flux (29, 42). Glycolysis (i.e., hepatic lactate release) increased maximally without an increase in F_2,6_P_2_ at 30 min, indicating that the ‘push’ from glycogenolysis was sufficient to drive flux through the glycolytic pathway. This suggests that substrate mass was a more important regulatory factor than enzyme activity at this stage of the response. By 4h, glucagon limited hyperglycemia-induced F_2,6_P_2_ production, and this probably explains the fall in hepatic lactate release despite hyperglycemia.

The effects of glucagon on glucose production characteristically waned with time. This was due to a decrease in net hepatic glycogenolysis rather than changes in hepatic glycolytic or gluconeogenic fluxes. Several possible reasons for glucagon’s evanescence have been suggested, including opposition by hyperglycemia and hyperinsulinemia, a depletion of hepatic glycogen, a decline in the amount of hepatic cAMP due to adenylate cyclase inhibition or phosphodiesterase activation, a desensitization of hepatic glucagon receptors, and/or CNS mediated suppression of HGP by glucagon (16). A transient increase in cAMP has been considered a leading explanation since glucagon enhances net splanchnic plasma cAMP release with a spike-decline pattern (11, 31). On the other hand, studies performed primarily in isolated rat hepatocytes have led to the notion that cAMP cannot mediate the physiologic effects of glucagon since pharmacologic levels (>350 pg/ml) of the hormone were required for adenylate cyclase activation in those studies (49). The present study refutes this concept since cAMP was markedly elevated by the physiologic rise in glucagon. Of note, we found that although cAMP decreased from its 30 min peak value, its levels remained elevated 4-fold even after 4h. Although GP activity was slightly lower at 4h vs 30 min, this was matched by a similar change in GS activity, such that over time there was no decline in the GP/GS ratio. Desensitization of the glucagon receptor or loss of downstream glucagon signaling provide unsatisfactory explanations since the covalent effects of glucagon on GS (inhibitory) and GP (stimulatory) activities were largely sustained over time, as was the sustained increase in hepatic alanine extraction. Depletion of liver glycogen is also an unlikely cause of the loss of glucagon action since glucagon only reduced glycogen content by about 20% at 4h. Recent rodent studies suggested that negative feedback by hypothalamic glucagon signaling can inhibit glucose production. When glucagon was administered directly into the brain, in those studies, HGP was suppressed, but when activation of the hypothalamic glucagon receptor or PKA were prevented the effect was blocked (1, 37). On the other hand, we found in dogs that while physiologic engagement of the brain by glucagon altered hepatic gluconeogenic / glycogenolytic carbon flux, it was not responsible for the transient fall in HGP following the stimulation of HGP during a rise in glucagon (17).

Hyperglycemia most likely contributed to the waning of glucagon’s effect on glycogen breakdown over time. Indeed, glucose inhibits glycogen phosphorylase after a latency which represents the time required to inactivate GP below the inhibitory threshold level (7). However, previous studies in the dog showed that when hyperglucagonemia was accompanied by an increase in plasma glucose, with insulin clamped at baseline, there was an 85% decrease in glucose production over 2h (10). On the other hand, glucose production only declined by 50% when the same rise in glucagon occurred if hyperglycemia was already present due to glucose infusion. Likewise, the effects of glucagon were more persistent in humans if changes in plasma glucose were prevented, but glucagon mediated stimulation of hepatic glucose production was still transient (20). Therefore, in addition to hyperglycemia, another component must contribute to the waning of HGP.

This factor is most likely G6P, which is inhibitory to net hepatic glycogenolysis (2, 3, 7), and which glucagon markedly increased. Hepatic G6P would have accumulated due to cAMP mediated activation of GP and the fall in glycolytic flux over time. Additionally, G6P increases glucokinase activity by promoting the enzyme’s translocation from the nucleus to the cytosol, which would increase hepatic glucose uptake and thus help maintain elevated G6P levels (2). G6P did not accumulate in the hyperglycemic control group because hyperglycemia would have suppressed GP and there was a sustained increase in glycolytic flux. G6P allosterically activates GS, which is coupled to an inactivation of GP, thus G6P opposes the effect of glucagon on net hepatic glycogenolysis (2, 3, 7). Therefore, it seems likely that increased hepatic G6P and glucose levels caused allosteric regulation of the glycogen-metabolizing enzymes and glucokinase that overcame the initial effects of cAMP. In contrast, G6P has no effect on hepatic amino acid uptake, so it is understandable that there was no time dependent loss of glucagon’s effect on that process. Thus, the present data suggest that the waning effect of glucagon on HGP seen in the absence of a rise in insulin may be due to a secondary increase in hepatic G6P and glucose levels, as opposed to tachyphylaxis of glucagon’s effect on cAMP concentration or the subsequent phosphorylation state of the rate-limiting enzymes involved in glycogen metabolism.

Neither hyperglycemia nor hyperglucagonemia modified acute (<4h) hepatic gluconeogenic flux. Glucagon’s inability to affect gluconeogenic flux is due in part to its lack of effect on gluconeogenic precursor supply from non-hepatic tissues. This process is highly dependent on substrate supply and NEFA availability (oxidized to support gluconeogenic flux) (19), but the glucagon receptor is not expressed in muscle, is minimally present in adipose tissue, and is not known to increase precursor flux to the liver (21, 26, 34). Indeed, in the present study, elevated glucagon did not bring about net hepatic lactate uptake at any point, nor did it augment the delivery of glycerol, alanine, or NEFA to the liver. Although glucagon stimulated hepatic amino acid transport by enhancing amino acid fractional extraction, the effect was offset by a decrease in blood amino acid levels. Thus, net hepatic amino acid uptake was maintained at a near-basal rate.

Glucagon levels typically increase to protect against low glucose, whereas glucagon was studied in the presence of hyperglycemia in the present study. This was done to avoid the complicating effects of hyperinsulinemia and other counterregulatory hormones. Several studies have indicated that elevations in glucagon during settings such as fasting, exercise, and hypoglycemia can enhance AMPK Thr172 phosphorylation and activity (6, 24, 33, 45). Conversely, in the presence of hyperglycemia hyperglucagonemia was only able to increase P-Thr172 AMPK at 30 min in the present study, indicating that some of glucagon’s effects are context dependent. It has been suggested that glucagon’s effects on AMPK are mediated by cellular ATP depletion caused by increased gluconeogenesis (32) but gluconeogenesis was not altered, which may explain why glucagon had no effect on AMPK. It is worth noting that hyperglycemia (regardless of the glucagon concentration) increased AMPK Ser485 phosphorylation. We previously observed that this site was phosphorylated during hyperinsulinemia and was correlated with decreased Thr172 phosphorylation (45). Thus, phosphorylation at Ser485 may modify glucagon’s ability to phosphorylate AMPK’s Thr172 site.

While recent studies have characterized the impact of acute and chronic glucagon receptor agonism and/or antagonism on liver gene expression in rodents (14, 18, 54), our study is unique in that we examined the acute transcriptional impact of hyperglucagonemia in a dog model under conditions in which the ensuing effects on blood glucose and insulin were controlled for and matched across groups. This experimental design permitted discrimination of the effects of hyperglucagonemia and hyperglycemia per se (i.e., GGN+GLU group) from those of basal glucagon and hyperglycemia (i.e., GLU group) on liver gene expression. Interestingly, the most prominent hepatic transcriptomic signature in the GGN+GLU group was broad activation of genes encoding proteins involved in hepatic mitochondrial oxidative phosphorylation (OXPHOS). Stimulation of respiration in rat liver mitochondria isolated from glucagon pre-treated animals was first described by Yamazaki and colleagues nearly 50 years ago (25, 65). More recently, Perry et al. (41) showed that glucagon stimulated hepatic mitochondrial oxidation in mice in both acute and chronic contexts in a manner requiring the mitochondrial-associated inositol triphosphate receptor 1 (41). Our findings support and extend these earlier observations by demonstrating that acute physiologic hyperglucagonemia and concomitant hyperglycemia – when compared to matched hyperglycemic controls – uniquely activates an OXPHOS gene expression program that presumably supports the heightened energy demands placed on the liver to drive ATP-consuming processes like gluconeogenesis and ureagenesis. The coupling of glucagon action with induction of genes that promote mitochondrial energy production seemingly conflicts with the findings of Berglund et al. (6) who showed that a physiologic increase in glucagon promotes hepatic energy depletion in mice, as evidenced by marked reduction in the liver ATP/AMP ratio and activation of AMPK. However, Berglund and colleagues (6) employed a unique hyperglucagonemic-euglycemic clamp protocol to isolate hepatic glucagon action in the absence of hyperglycemia or hyperinsulinemia, whereas we examined hepatic glucagon action in a setting of basal insulin and hyperglycemia. While we did not measure hepatic adenine nucleotide levels in our study, our gene expression findings suggest that the glycemic level differentially influenced the impact of glucagon on hepatic energy state. Alternatively, OXPHOS gene activation after 4h of hyperglucagonemia may reflect a compensatory response to a prior fall in the ATP/AMP ratio, though such a fall was not reflected in the phosphorylation state of AMPK at 30min or 4h. Future studies are needed to clarify the temporal impact of hepatic glucagon action on mitochondrial oxidation and hepatic energy state under carefully controlled physiological conditions. Such studies will be of great interest as chronic glucagon infusion or glucagon receptor agonism in diet-induced obese rodents was shown to reverse hepatic steatosis via stimulation of hepatic mitochondrial fatty acid oxidation (28, 41).

Unexpectedly, genes annotated to pathways involved in skeletal muscle development (i.e., myogenesis) and wound healing, fibrosis, and metastasis (i.e., EMT) were highly enriched among the downregulated genes in the GGN+GLU group. Since myogenesis, EMT, and angiogenesis (another pathway significantly downregulated in the GGN+GLU group) are categorized together as development-related processes (30), our data suggest that glucagon signaling may reinforce mature hepatocyte identity by reducing the expression of genes involved in developmental pathways. Furthermore, functional GO terms associated with these pathways highlighted several biological processes involved in ECM assembly and organization, collagen biosynthesis and degradation, and external encapsulating structure organization. The ECM comprises a heterogeneous network of proteoglycans, polysaccharides, and proteins that provide a structural scaffold for cells and tissues, transduce signals through ECM-receptor interactions, and undergo dynamic changes in their abundance and composition to maintain tissue homeostasis in response to injury (40, 64). Although links between obesity-induced ECM remodeling and impaired insulin action in the liver, skeletal muscle, and adipose tissue have been established (40, 64), few if any studies have examined the impact of ECM remodeling on glucagon action or vice versa. Here, we show that hyperglucagonemia leads to downregulation of numerous matrisome genes encoding various collagens and ECM-related proteins. How glucagon signaling may regulate these processes and whether they manifest as alterations in the abundance and/or composition hepatic ECM remains to be determined.

In summary, glucagon acutely stimulates a rapid and profound increase in HGP. This is due exclusively to increased net hepatic glycogenolysis, which is associated with increased cAMP formation and coordinated activation of GP and inhibition of GS. The waning effect of glucagon on hepatic glucose metabolism occurred even though cAMP remained substantially elevated and the effects on glycogen-metabolizing enzymes were sustained. Hyperglucagonemia caused hyperglycemia and a substantial increase in hepatic G6P, and the allosteric effects of glucose and of G6P on GS, GP, and glucokinase are likely to have mediated the evanescence of glucagon action. Hyperglucagonemia also induced gene expression programs associated with upregulation of hepatic mitochondrial oxidation and metabolism, and downregulated genes involved in ECM organization and developmental processes. These results emphasize glucagon’s potent effects on glucose metabolism, while revealing novel insights into hepatic glucagon signaling that are relevant to the evaluation of glucagon antagonists as therapeutic options in diabetes treatment.

## Data Availability

Data will be made available upon reasonable request.

## Supplemental Data

Supplemental Table 1 (representative Western blots for Figure 4) is available at the following DOI: https://doi.org/10.17632/rjhncsmp58.1

## Grants

This research was supported by National Institutes of Health Grant DK18243 and the Diabetes Research and Training Center Grant SP-60-AM20593. Hormone analysis was performed by Vanderbilt’s Hormone Assay and Analytical Services Core, and surgical and experimental expertise was provided by Vanderbilt’s Large Animal Core, both of which are supported by the Vanderbilt University Medical Center Diabetes Research and Training Center grant DK20593. CJR was supported by the American Diabetes Association Mentor-based Fellowship. ADC was supported by the Vanderbilt Jacquelyn A. Turner and Dr. Dorothy J. Turner Chair in Diabetes Research.

## Author Contributions

CJR, ADC, and DSE conceived and designed research; CJR, GK, BF, and DSE performed experiments; KCC, CJR, MS, JW, JID, EPD, PJR, ADC, and DSE analyzed data, KCC, CJR, ADC, and DSE interpreted results of experiments; KCC, MS, ADC, and DSE prepared figures; KCC, CJR, ADC, and DSE drafted manuscript; KCC, CJR, JW, GK, JID, DSE and ADC edited and revised manuscript; KCC, CJR, MS, JW, GK, JID, BF, EPD, ADC, and DSE approved final version of manuscript.

### Author Notes

• Correspondence: D. Edgerton

## Supplemental figure legends

**Fig. S1.**
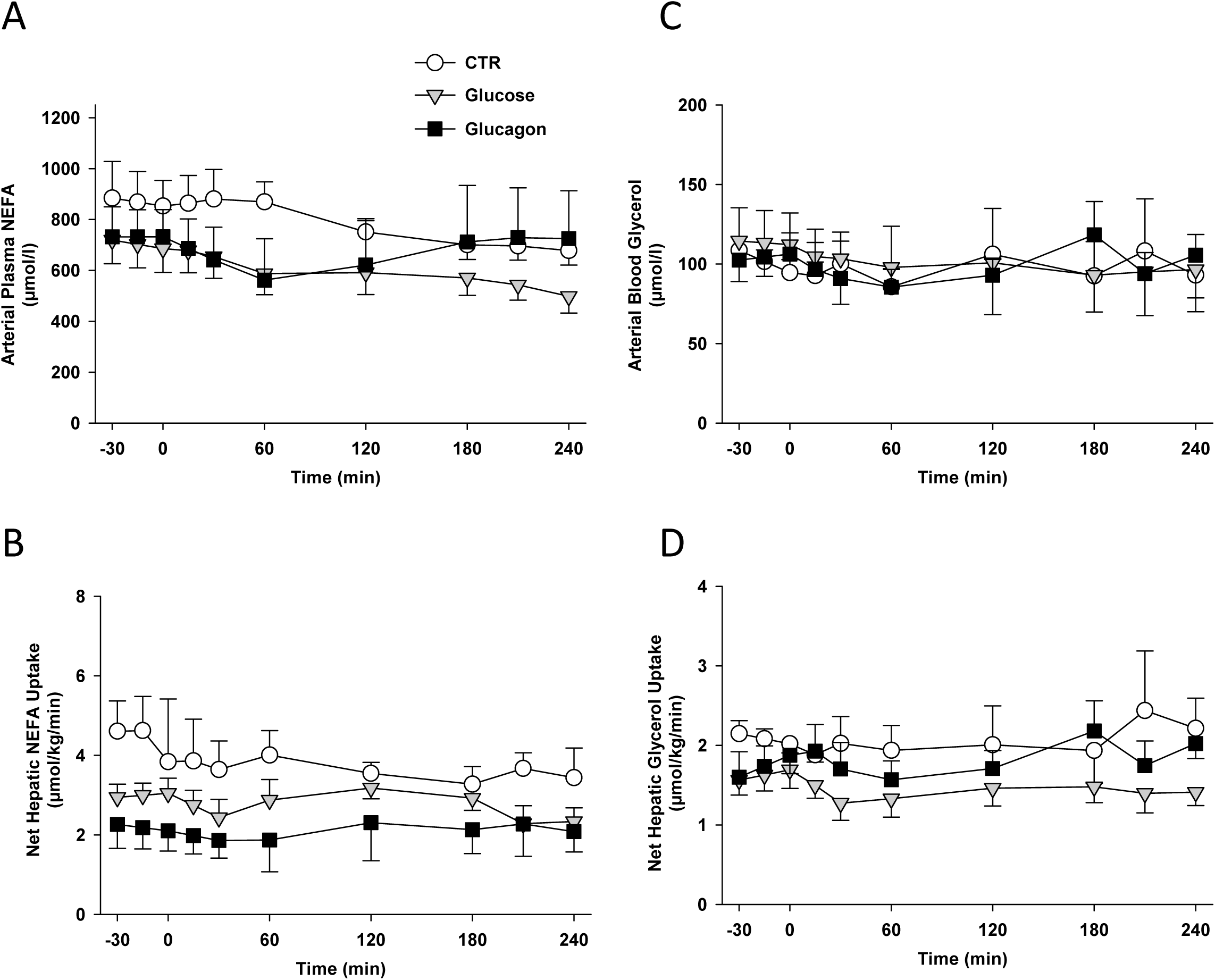
Arterial plasma or blood (A) NEFA levels, (B) net hepatic NEFA uptake, (C) glycerol levels and (D) net hepatic glycerol uptake in 18h fasted conscious dogs during the basal (–30 to 0 min) and experimental (0-240 min) periods. Data are means ± SEM, in the CTR (n=6 from –30 to 240 min), GLU (n=8 from –30 to 30 min; n=4 from 30 to 240 min), and GGN+GLU (n=8 from –30 to 30 min; n=4 from 30 to 240 min) groups.

**Fig. S2.**
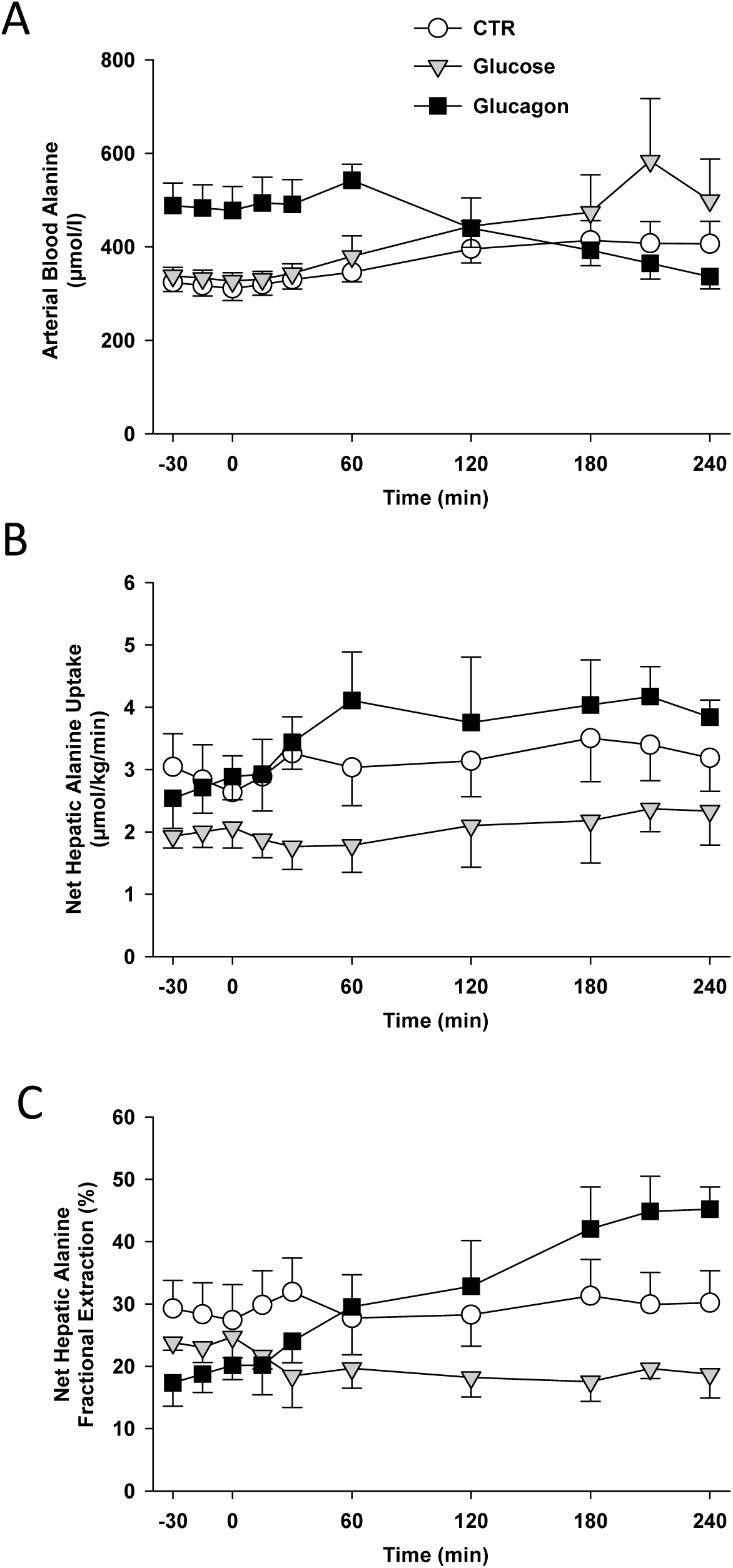
(A) Arterial blood alanine levels, (B) net hepatic alanine uptake, and (C) net hepatic alanine fractional extraction in 18h fasted conscious dogs during the basal (–30 to 0 min) and experimental (0-240 min) periods. Data are means ± SEM, in the CTR (n=6 from –30 to 240 min), GLU (n=8 from –30 to 30 min; n=4 from 30 to 240 min), and GGN+GLU (n=8 from –30 to 30 min; n=4 from 30 to 240 min) groups. Net hepatic alanine fractional extraction was greater (*P* <0.05) in the GGN+GLU group compared to the GLU group between 180-240 min.

**Fig. S3.**
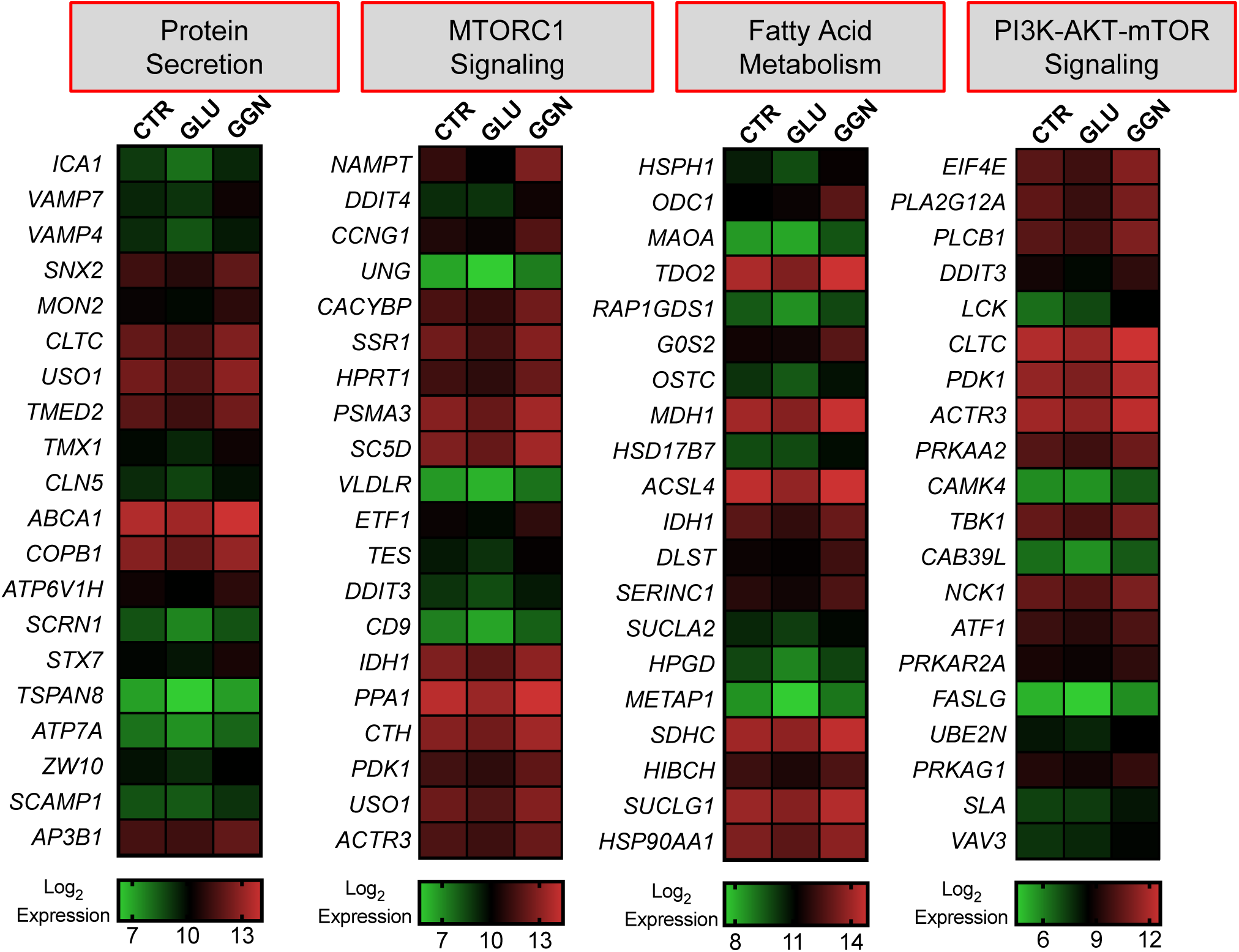
Heat map representation of the top 20 leading-edge transcripts underlying the glucagon-regulated positive enrichment signals for protein secretion, MTORC1 signaling, fatty acid metabolism, and PI3K-AKT-mTOR signaling as identified in Fig. 6B. Each row represents a different glucagon-regulated gene, whereas each column represents the mean log_2_ normalized expression value for these genes in the CTR, GLU, and GGN+GLU groups (n=3 each). Expression values are represented as colors and range from green (lowest expression) to red (highest expression).

---

**Supplemental Table 1.**
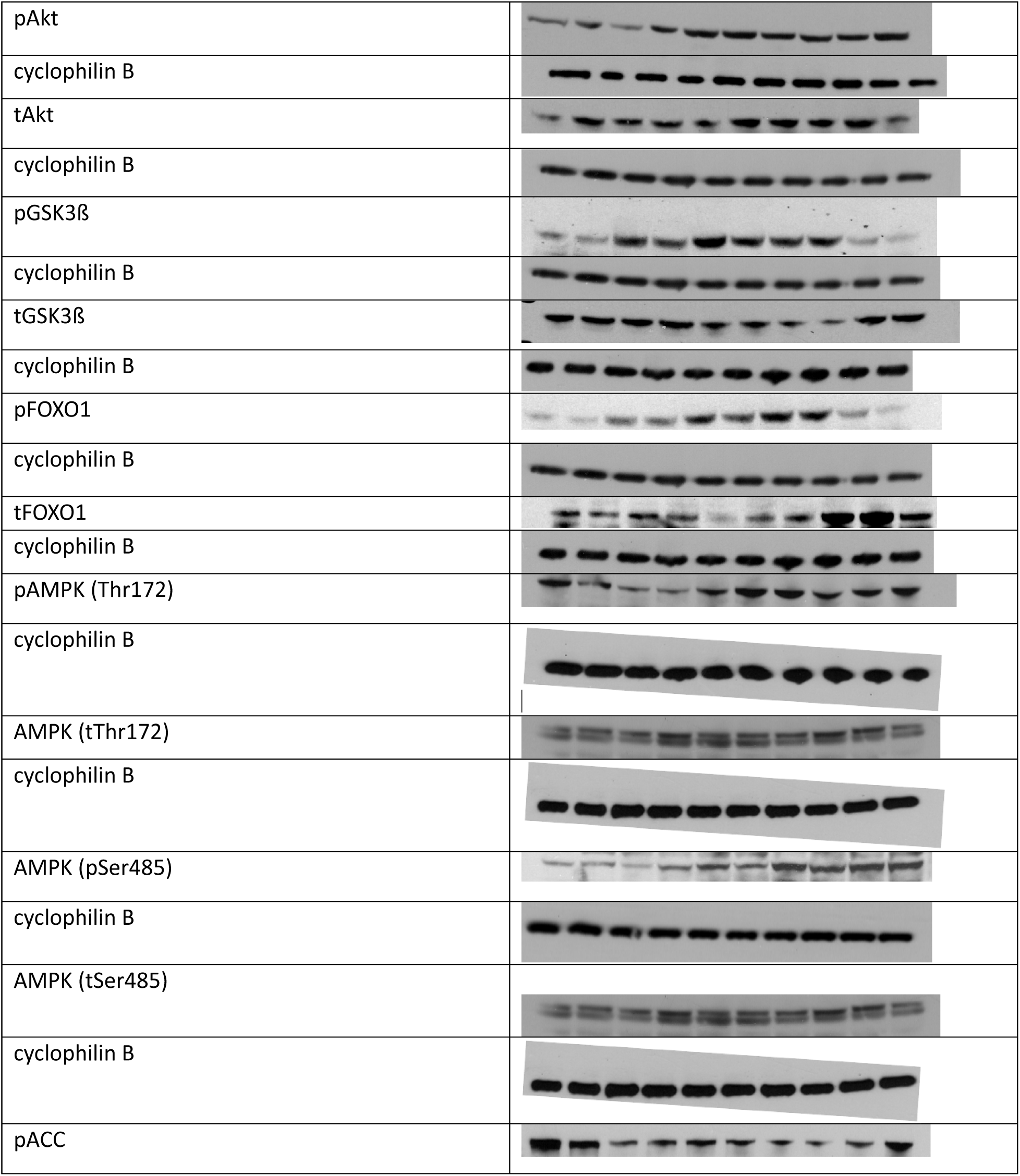
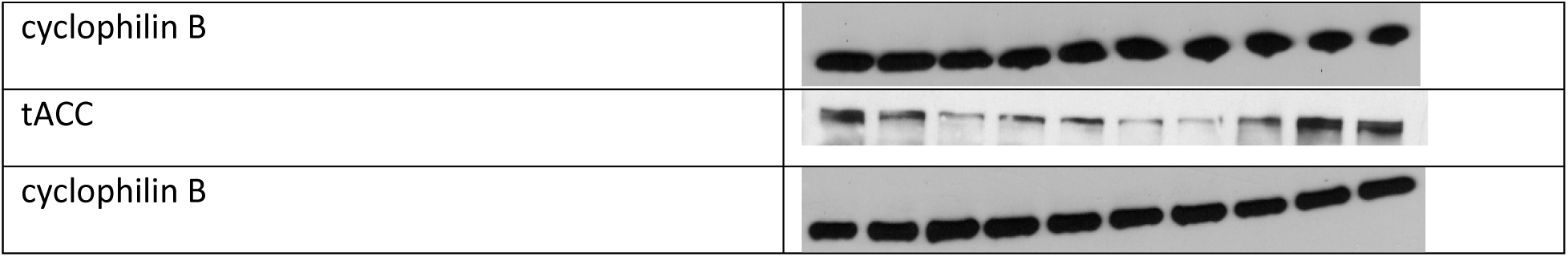
Representative Western blots. Phosphorylated and total proteins, with cyclophilin B from each respective gel. Lanes 1-2: CTR, 3-4: GLU 30 min, 5-6: GGN+GLU 30 min, 7-8: GLU 4h, 9-10: GGN+GLU 4h

